# ATP11A and ATP11C are plasma membrane phosphatidylserine flippases in *in vitro* human megakaryocytes

**DOI:** 10.64898/2026.01.30.702765

**Authors:** Ruby Baxter, Alexi Crosby, Holly R. Foster, Winnie Lau, Amie K. Waller, Cedric Ghevaert, Matthew T. Harper

## Abstract

Thrombotic diseases are the major worldwide cause of cardiovascular death. Platelets prevent blood loss following injury (haemostasis), but inappropriate and excessive platelet activation can lead to thrombosis. Platelet activation must be tightly controlled. Pro-coagulant platelets expose phosphatidylserine (PS), enabling coagulation complex assembly, enhancing thrombin generation and thrombosis. PS is normally restricted to the inner leaflet of the plasma membrane by ‘flippase’ (aminophospholipid translocase) activity. However, the flippase protein(s) responsible for this crucial activity in platelets remains unidentified. The P4 ATPases ATP11A and ATP11C, regulated by their obligatory partner CDC50A, flip PS at the plasma membrane in a range of different cell types. To investigate platelet flippases, human induced pluripotent stem cells (hiPSCs) were forward-programmed into CD41^+^/CD42^+^ megakaryocytes, the platelet precursor. Wildtype (WT) forward-programmed megakaryocytes showed similar flippase activity to human platelets with internalisation of NBD-PS that could be inhibited by high cytosolic Ca^2+^ or *N*-ethylmaleimide (NEM). We then generated CDC50A, ATP11A or ATP11C single knockout and ATP11A/11C double knockout (DKO) hiPSCs using CRISPR-Cas9. CDC50A-KO, ATP11A-KO, ATP11C-KO and DKO hiPSC clones successfully formed CD41a^+^/CD42a^+^ mature megakaryocytes. CDC50A-KO megakaryocytes bound Annexin V when unstimulated and had no remaining NEM-sensitive flippase activity indicating the involvement of a P4-ATPase. Although ATP11A-KO and ATP11C-KO megakaryocytes had similar flippase activity to WT clones, DKO clones had inhibited NBD-PS internalisation compared to WT and had no remaining NEM-sensitive flippase activity. This indicates that the CDC50A-regulated P4-ATPases ATP11A and ATP11C act together at the megakaryocyte plasma membrane and are responsible for PS flippase activity and therefore likely responsible in human platelets.

## Introduction

Circulating platelets prevent blood loss following injury (haemostasis), but inappropriate and excessive platelet activation can lead to thrombosis. Platelet activation must be tightly controlled to prevent thrombotic disease, the major worldwide cause of cardiovascular death^1^. Conversely, failure of normal haemostasis is also a major cause of preventable deaths. Mortality following traumatic haemorrhage, for example, remains high despite major advances in immediate care^2^. Post-partum haemorrhage is the primary cause of maternal mortality worldwide^3^. Phosphatidylserine (PS) exposure by activated platelets plays a key role in thrombosis and haemostasis. This PS exposure enhances coagulation by facilitating the assembly of the tenase and prothrombinase complexes, increasing local thrombin generation and potentiating thrombosis. Despite this, the molecular regulation of platelet PS exposure remains incompletely described.

PS is normally restricted to the inner leaflet of the plasma membrane of unactivated platelets. Within a growing thrombus, a sub-population of activated platelets, known as ‘pro-coagulant platelets’, expose PS in the outer leaflet^4^. PS exposure in pro-coagulant platelets requires activation of the phospholipid scramblase, TMEM16F^5^. This occurs downstream of a high cytosolic Ca^2+^ signal following mitochondrial permeability transition pore opening^7,8^. The contribution of PS exposure to thrombosis and haemostasis is clearly demonstrated by a moderate bleeding risk in patients with Scott Syndrome (with mutations in TMEM16F^9^), and reduced thrombosis in *Tmem16f*^*-/-*^ mice^10,11,12^. PS exposure must therefore be tightly controlled to prevent inappropriate PS exposure that would increase risk of thrombosis while rapidly providing PS exposure when needed for haemostasis.

Membrane asymmetry is maintained in unstimulated (and activated but non-coagulant) platelets by phospholipid ‘flippase’ activity that mediates inward translocation of PS. This flippase activity requires ATP and is specific for PS and phosphatidylethanolamine (PE) over phosphatidylcholine (PC)^13^. The activity is inhibited by the alkylating agent, *N*-ethylmaleimide (NEM)^14^, and by the high cytosolic Ca^2+^ concentration in pro-coagulant platelets^6^. If flippase activity is maintained in pro-coagulant platelets, overall PS exposure and platelet-dependent thrombin generation are reduced, making this a potential anti-thrombotic approach^15^. Despite PS flippase activity being first reported in the late 1980s^16^, the proteins responsible in platelets are currently unidentified.

Flippase activity in many cell types is due to P4 ATPases. P4 ATPases are known to be inhibited by high cytosolic Ca^2+^, NEM and vanadate^17^. There are 14 P4 ATPases in the human proteome^18^, with most P4 ATPases requiring partnership with CDC50A for correct cellular localisation^19,20^ and function. Mutations in CDC50A result in loss of membrane asymmetry demonstrated by Annexin V binding^21,22,23^. The P4 ATPases ATP11A and ATP11C are known to be present at the plasma membrane^24,25,26^ and can contribute to PS translocation in murine W3 cells ^23,27,28^ and human BeWo cells^29^. Whilst the identity of the platelet PS flippase is unknown, the P4 ATPase ATP11C has been identified as the major flippase protein in human erythrocytes, with several cases of haemolytic anaemia due to ATP11C mutations. These ATP11C mutations resulted in a loss of PS translocation at the plasma membrane of erythrocytes ^30,31,32,33,34^. In one case, platelet count was measured at 125-149x10^9^/L (normal range 150-450x10^9^/L) but flippase activity was not tested^31^. Identification of the protein responsible for PS translocation at the plasma membrane of human platelets is complicated by their lack of a nucleus, preventing gene editing or siRNA approaches.

In this study, we use a novel approach to identify the protein(s) responsible for the inward translocation of PS at the plasma membrane of megakaryocytes to identify the most likely candidates for the platelet flippase protein. Mutations were introduced into candidate flippase proteins (CDC50A, ATP11A and ATP11C) in human induced pluripotent stem cells (hiPSCs) using CRISPR Cas9. Knockout hiPSCs were used to generate megakaryocytes *in vitro* using Forward Programming (FoP) technology^35,36^. Flippase activity was measured in CDC50A-KO, ATP11A-KO, ATP11C-KO and ATP11A/ATP11C double knockout (DKO) megakaryocytes, identifying flippase activity as CDC50A dependent and both ATP11A and ATP11C as flippases jointly responsible for PS internalisation at the plasma membrane of FoP megakaryocytes. ATP11A and ATP11C are likely to be responsible for flippase activity in human platelets.

## Methods

### Human blood preparation

Blood collection was performed with approval from the Human Biology Research Ethics Committee, University of Cambridge in accordance with the Declaration of Helsinki. Blood was collected from healthy, drug free volunteers into 3.2% sodium citrate vacutainers. Platelet rich plasma (PRP) was separated (centrifugation: 200g, 10 minutes) following addition of Acid citrate dextrose (ACD: 97mM sodium citrate, 62.5mM citric acid, 111mM D-glucose). PRP was diluted 1:1 with N-2-hydroxyethylpiperazine-N’-2-theanesulfonic acid (HEPES)-buffered saline (HBS: 10mM HEPES free acid, 135mM NaCl, 3 mM KCl, 0.34 mM NaH_2_PO_4_, 1mM MgCl_2_.6H_2_0, pH 7.4, 5mM D-glucose) and supplemented with PGE_1_ (100nM) and apyrase (0.02 units/ml) before pelleting platelets by centrifugation (600g, 10 minutes). Platelets were resuspended in HBS at a density of 5x10^7^/ml. Following the removal of PRP, the erythrocytes were diluted 1:1 with HBS and washed (centrifugation: 1000g, 10 minutes). This was completed three times to remove plasma. Erythrocytes were diluted in HBS to a density of 5x10^7^/ml.

### Generation of knockout hiPSCs

Human inducible hiPSC lines iQOLG1.1A (parental line HPSI1113i-qolg_1: HipSci) and iLIPSC-GR1.1.11A (parental line LIPSC-GR1.1: proprietary) were generated by the Ghevaert Group. iPSCs were maintained in culture on recombinant vitronectin (Thermo Fisher Scientific; 0.5μg/cm^2^) in mTeSR Plus media (StemCell Tech) and passaged using TrypLE Select (Glibco). iPSCs were dissociated using TrypLE and nucleoporated (Lonza 4D-Nucleofector) to introduce gRNAs (Synthego; 183pmol) and Cas9 enzyme (Integrated DNA Technologies; 10μg). We used the following single gRNAs. For CDC50A: 5’ – AUAGCUCCACAAGGAGCAAU. For ATP11A: 5’ – GGACAGCAGGACCAUCUACG. For ATP11C: 5’ – UCACUACAGCCAGUCUUGAU.

Edited populations were single cell live/dead sorted into clones which were expanded for DNA extraction (KAPA Express Extract) and PCR-amplified (KAPA Robust Hotstart PCR Kit) using the following conditions: 3-minute hot start at 95°C followed by 40 cycles of a denaturation step at 95°C for 15 seconds and an annealing and amplification step at 72°C for 1 minute. PCR products were genotyped by Sanger sequencing (Source Bioscience) and analysed using Inference of CRISPR Edits (ICE). We used the following primers (Merck). For CDC50A: 5’ – ACAGACCTGTGTATTATAAGTGAGA and 5’ – ACCCCATTCAATACTCGATTTAGTC. ATP11A: 5’ – CAGCCCTGAACGATGCTCT and 5’ – CCTGATCCGAAGCCCTTGC. For ATP11C: 5’ – ATGCTGAGATACTCTCAAAGCCC and 5’ – GGTGATGGGAGGCAGTATTGT.

### Forward programming (FoP)

Inducible FoP was carried out as previously described^36^ with doxycycline (iQOLG1.1A: 0.0325μg/ml, iLIPSC-GR1.1.11A: 0.02μg/ml) used to induce the expression of GATA1, FLI1 and TAL1 using a TET-ON system. Briefly, 24 hours prior to FoP, cells were dissociated using TrypLE and seeded on vitronectin at 100,000 cells/ well of a 6 well plate. Following this, there was a 2-day induction of mesoderm (FGF2; in house; 20ng/ml, BMP4; Qkine; 10ng/ml) with 24-hour treatment with Chiron (30nM) followed by megakaryocyte specific media (STable. 1) for 18 days containing thrombopoietin (Peprotech; 20ng/ml) and Stem Cell Factor (Thermofisher; 25ng/ml). Cells were fed every 2-3 days. Cells were assessed for megakaryocyte phenotype at Day 20 using specific antibodies for CD41a (BD Biosciences: 561422) and CD42a (Miltenyi Biotech: 130-123-831).

### Assessment of flippase activity

Platelets and FoP megakaryocytes were stained with PE-conjugated Annexin V to detect PS exposure in resting and stimulated cells (A23187: 10μM, 2mM CaCl_2_, 10 minutes).

NBD (7-nitro-2-1,3-benzoxadiazol-4-yl)-labelled phosphatidylserine (NBD-PS) was used to assess translocation of PS at the plasma membrane. Platelets and megakaryocytes were treated with *N*-ethylmaleimide (NEM; 5mM, 15 minutes), A23187 (10μM, with 2mM CaCl_2_, 10 minutes) or sodium orthovanadate (vanadate; 0.1mM/10mM, 15 minutes) prior to the addition of 5μM NBD-PS and incubation at 37°C. At various time points, cells were sampled into HBS±2% bovine serum albumin (BSA). BSA extracts NBD-PS in the outer leaflet. The percentage non-extractable NBD-PS fluorescence was calculated by dividing the median fluorescence intensity in the HBS sample by that in the BSA-treated sample.

Annexin V and NBD-PS fluorescence were measured using a BD Accuri C6 flow cytometer. 10000 platelet events were collected, and 1000 viable FoP megakaryocyte events were collected, both identified by their forward and side scatter profile.

## Data analysis and statistics

Data presented as mean ± standard error of the mean (SEM). Biological replicates for human platelets and erythrocytes represent an independent blood preparation from different donors. Biological replicates for *in vitro* megakaryocytes represent cells from a different passage number that have independently undergone a forward programming protocol. The number of biological repeats is given in the figure legend. Statistical tests used are described in the figure legends. Time courses were fitted to logistic curves^37^ .

## Results and Discussion

We use megakaryocytes differentiated from iQOLG1.1A hiPSCs by forward programming (FoP) as a genetically tractable model of human platelets. These FoP megakaryocytes expressed platelet/megakaryocyte-specific markers CD41a and CD42a. To validate FoP megakaryocytes as a model for platelet flippase activity, PS exposure was measured by Annexin V binding, and flippase activity was measured by NBD-PS internalisation (Fig. 1A). PS exposure was very low in unactivated platelets and FoP megakaryocytes. Stimulation with the calcium ionophore, A23187, in the presence of 2mM extracellular CaCl_2_, resulted in PS exposure in most platelets and FoP megakaryocytes, as expected (Fig. 1B).

**Figure 1:**
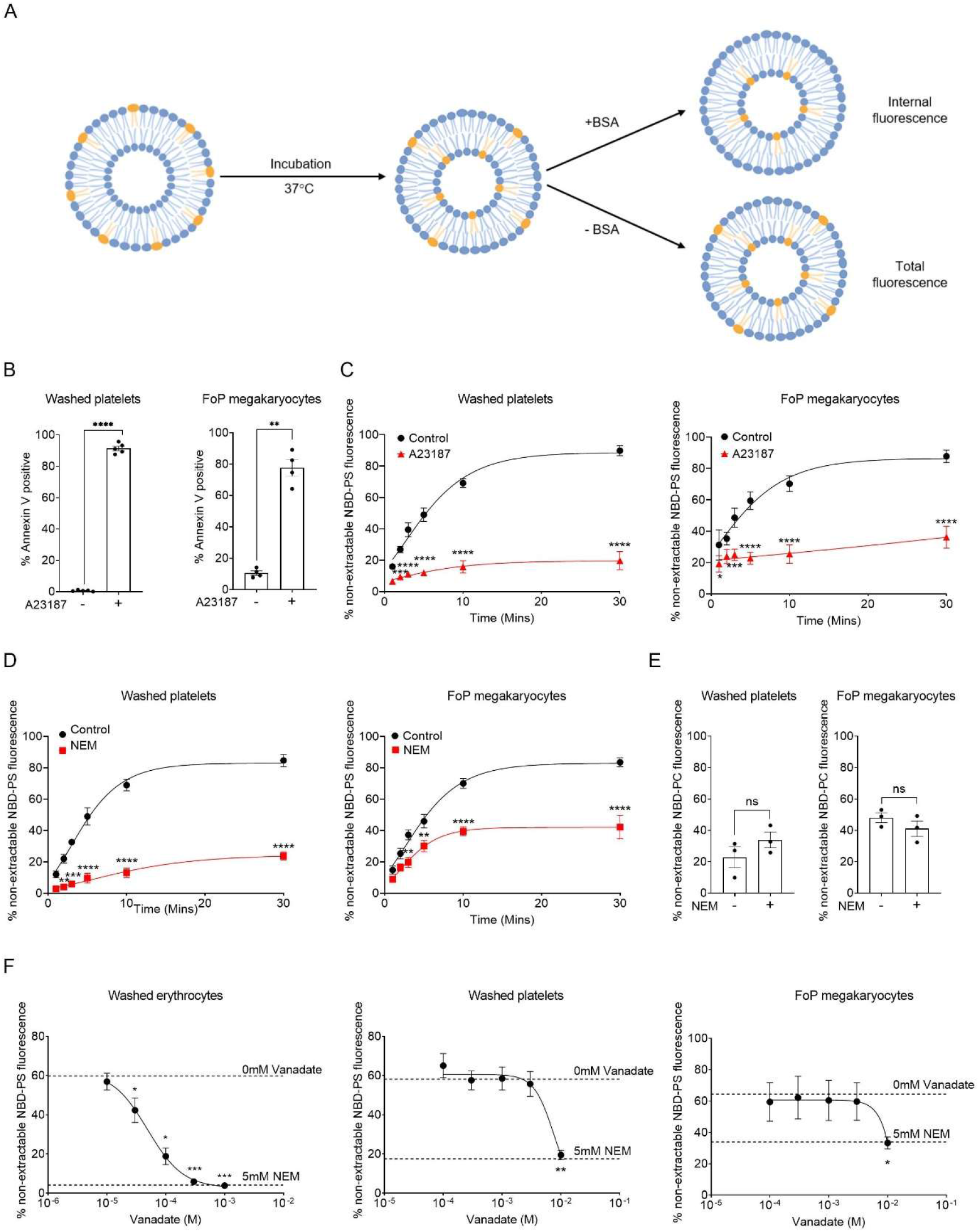
Flippase activity in FoP megakaryocytes is similar to that in washed human platelets. **A**. Schematic diagram of NBD-PS flippase assay. BSA extracts NBD-PS from the outer leaflet. **B**. Washed human platelets (5x10^7^/ml) and FoP megakaryocytes from iQOLG1.1A iPSCs (5x10^5^/ml) were stimulated with A23187 (10μM: 10 minutes; with 2mM CaCl_2_). Cells were then stained with Annexin V to detect PS exposure. **C-F**. Human platelets or FoP megakaryocytes were treated as indicated with A23187, NEM (5μM: 15 minutes), vanadate (various concentrations: 15 minutes), or vehicle controls, and incubated with NBD-PS (**C**,**D**,**F** 5μM: 37°C) or NBD-PC (**E** 5μM: 37°C: 30 minutes) for the indicated times before sampling into HBS± 2% BSA. **F**. Washed human erythrocytes (5x10^7^/ml) were incubated with vanadate (various concentrations: 15 minutes) and incubated with NBD-PS (5μM: 37°C: 60 minutes) before sampling into HBS± 2% BSA. Data analysed by paired Student’s T test **(B&E)**, one-way ANOVA **(F)** or two-way ANOVA **(C&D)** followed by Dunnett’s multiple comparisons (*p<0.05, **p<0.01, ***p<0.001, ****p<0.0001). Mean ± SEM (N=3-5).

Platelets and FoP megakaryocytes rapidly internalised NBD-PS (Fig. 1C), indicating flippase activity. Platelet flippase activity has previously been demonstrated as being inhibited by high cytosolic Ca^2+^ concentration^6^, and the alkylating agent NEM^14^, a known non-selective P4 ATPase inhibitor^17^. In both cells, stimulation with the Ca^2+^ ionophore, A23187, significantly inhibited NBD-PS internalisation, indicating inhibition of flippase activity (Fig. 1C). PS flippase activity was also inhibited by NEM (Fig. 1D). In contrast, NBD-PC internalisation, which is slower than NBD-PS internalisation, was not inhibited by NEM (Fig. 1E). The residual NBD-PS internalisation in NEM treated FoP megakaryocytes is therefore different to platelet flippase activity with NBD-PC also being transported. This therefore likely represents a flippase independent, non-selective, route of lipid uptake.

Flippase activity in other cell types, such as human erythrocytes, has been demonstrated as inhibited by vanadate^38^, but this has not been reported in human platelets. Whilst erythrocyte PS flippase activity was significantly inhibited by vanadate in the low micromolar range, 10mM vanadate was required for significant inhibition of flippase activity in washed human platelets and FoP megakaryocytes (Fig. 1F). All findings were replicated in FoP megakaryocytes derived from a second hiPSC line (iLIPSC-GR1.1.11A; SFig. 1). This demonstrates that FoP megakaryocyte flippase activity is similar to in platelets and hence they can be used as a model for platelet flippase activity.

P4-ATPase PS internalisation in other cells is dependent on CDC50A^21,22,23^. Loss of NBD-PS internalisation in CDC50A knockout megakaryocytes would confirm the role of a P4 ATPase in this process. CRISPR-Cas9 was used to generate CDC50A genetic knockout hiPSCs (iQOLG1.1A) and FoP megakaryocytes. Two CDC50A-KO clones were selected with different frame shift mutations in the target sequence (STable. 2, SFig. 2). These were forward programmed in 3 independent experiments alongside a wildtype clone and the unedited (parental) population (Fig. 2A) generating CD41a^+^/CD42a^+^ mature megakaryocytes (SFig. 6). Genetic loss of CDC50A resulted in constitutive PS exposure as demonstrated by high Annexin V binding, indicating loss of membrane asymmetry (Fig. 2B-C). CDC50A-KO megakaryocytes also had inhibited flippase activity measured by NBD-PS internalisation (Fig. 2D). This phenotype was replicated in CDC50A-KO FoP megakaryocytes from the iLIPSC-GR-1.1.11A line (SFig. 7). This confirms that NBD-PS internalisation in FoP megakaryocytes is due to CDC50A-dependent P4-ATPase activity.

**Figure 2:**
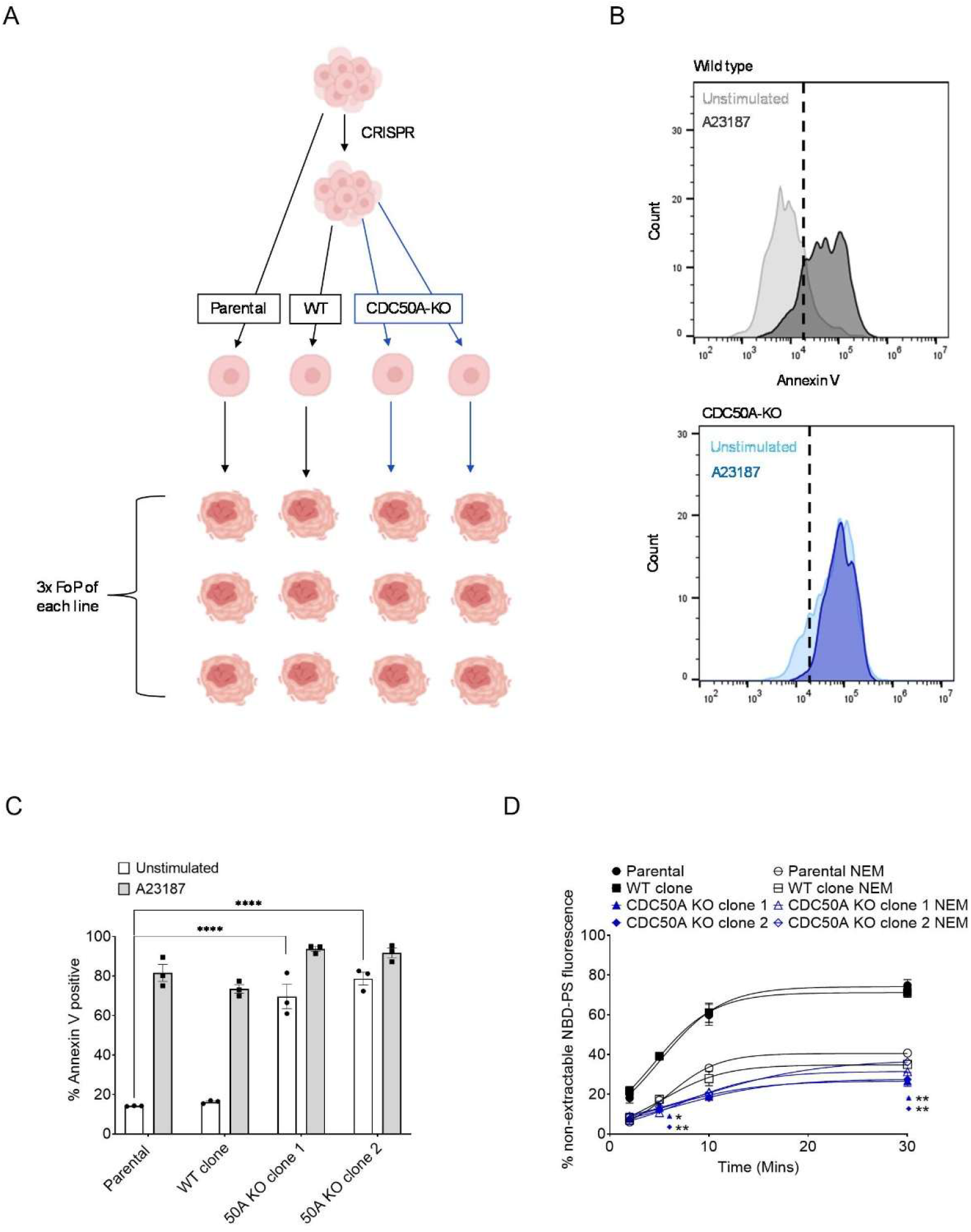
Flippase activity is lost in CDC50A-KO megakaryocytes. **A**. Schematic diagram of approach to phenotypic analysis. **B**. Representative histograms of Annexin V binding to unstimulated and A23187 treated WT and CDC50A-KO FoP megakaryocytes. **C**. CDC50A-KO FoP megakaryocytes from the iQOLG1.1A line were stimulated with A23187 (10μM: 10 minutes: with 2mM Ca^2+^) and stained with Annexin V to detect PS exposure. **D**. FoP megakaryocytes were incubated with NEM (5mM: 15 minutes) or vehicle control and incubated with NBD-PS (5μM: 37°C) before sampling into HBS± 2% BSA. Data analysed by two-way ANOVA followed by Dunnett’s multiple comparisons compared with untreated parental FoP MKs (*p<0.05, **p<0.01, ***p<0.001, ****p<0.0001). Mean ± SEM (N=3).

The difference in vanadate sensitivity between human erythrocytes and platelets could suggest a difference in the P4 ATPase responsible for PS translocation. This is supported by a patient with an ATP11C mutation having only very mild thrombocytopaenia but severe haemolytic anaemia^31^.

Hence, we initially considered ATP11A as a more likely candidate than ATP11C. ATP11A knockout (ATP11A-KO) hiPSCs were generated using CRISPR-Cas9 gene editing (STable. 3, SFig. 3). Flippase activity in two independent ATP11A-KO clones were compared with a WT clone isolated from the edited population and with the parental population which had not undergone gene editing. ATP11A-KO megakaryocytes had no constitutive PS exposure (Fig. 3A). The flippase activity was similar to that of WT megakaryocytes, and was inhibited by NEM (Fig. 3B) and a high cytosolic Ca^2+^ signal induced by A23187 (Fig. 3C). This demonstrates that despite loss of ATP11A, there was PS flippase activity remaining and hence ATP11A is not solely responsible for PS translocation. These findings were replicated in FoP megakaryocytes from the iLIPSC-GR-1.1.11A line (SFig. 7).

**Figure 3:**
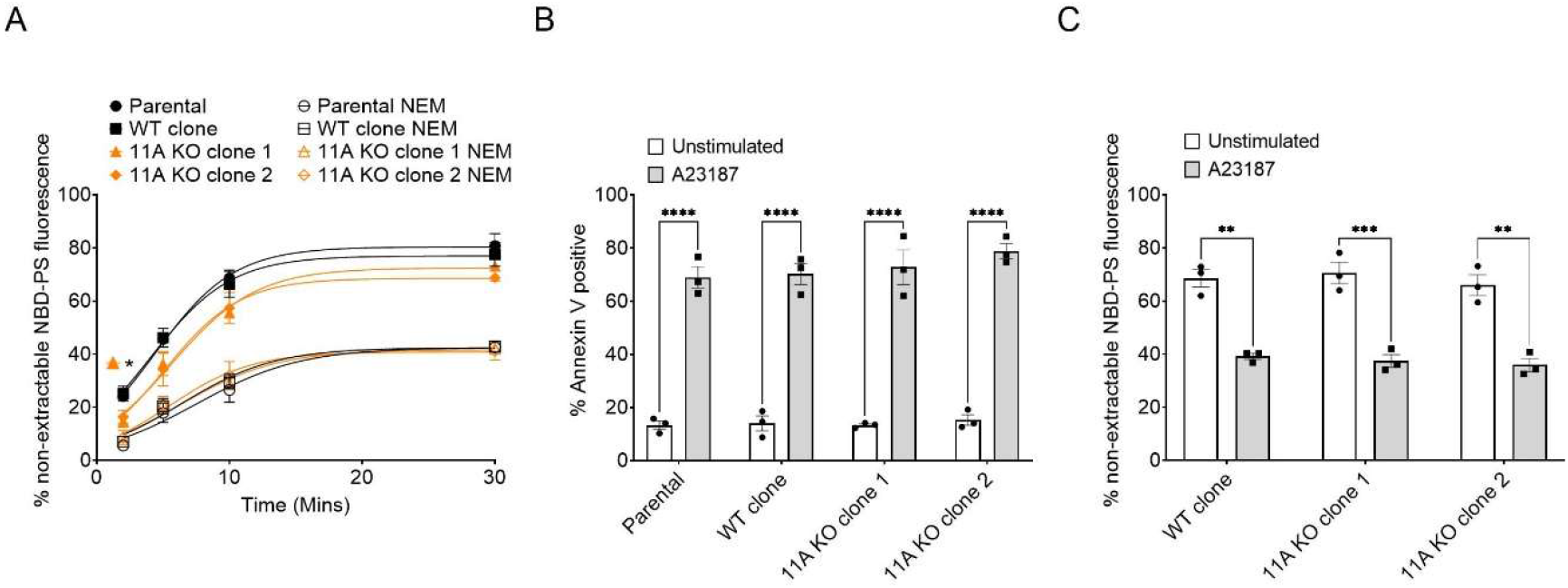
ATP11A-KO megakaryocytes have a phenotype similar to WT megakaryocytes. **A**. iQOLG1.1A FoP megakaryocytes were incubated with NEM (5mM: 15 minutes) or vehicle control and incubated with NBD-PS (5μM: 37°C) before sampling into HBS± 2% BSA. **B**. FoP megakaryocytes were stimulated with A23187 (10μM: 10 minutes: with 2mM Ca^2+^) and stained with Annexin V to detect PS exposure. **C**. FoP megakaryocytes were stimulated with A23187 (10μM: 10 minutes: with 2mM Ca^2+^) and incubated with NBD-PS (5μM: 37°C: 30 minutes) before sampling into HBS± 2% BSA. Data analysed by two-way ANOVA followed by Dunnett’s multiple comparisons compared with untreated parental FoP MKs (A) or unstimulated FoP MKs (B-C) (**p<0.01, ***p<0.001, ****p<0.0001). Mean ± SEM (N=3).

Consequently, we investigated the role of ATP11C in PS internalisation. ATP11C knockout (ATP11C-KO) and double knockout (DKO) hiPSCs were generated using simultaneous gene editing (STable. 3, SFig. 4-5). The ATP11C-KO and DKO hiPSCs successfully produced CD41a^+^/CD42a^+^ megakaryocytes (SFig. 6). Similarly to ATP11A-KO, ATP11C-KO megakaryocytes had no constitutive PS exposure, as measured by Annexin V binding (Fig. 4A) and had rapid NBD-PS internalisation that was inhibited by NEM (Fig. 4B) and A23187 treatment (Fig. 4C). However, when both ATP11A and ATP11C were deleted in the DKO megakaryocytes, there was full loss of NEM-sensitive NBD-PS internalisation (Figure 4D; replicated in the iLIPSC-GR-1.1.11A line: SFig. 7). PS flippase activity at the plasma membrane of these DKO megakaryocytes was therefore negligible, identifying ATP11A and ATP11C as acting together and redundantly to internalise plasma membrane PS into the inner leaflet. Despite low PS internalisation at the plasma membrane, there was no constitutive PS exposure as detected by Annexin V (Figure 4E). This suggests that plasma membrane lipid asymmetry is maintained even in the absence of rapid plasma membrane PS translocation. DKO megakaryocytes maintained their ability to expose PS following activation with A23187 (Figure 4E). This agrees with findings in ATP11A/ATP11C DKO mouse W3 cells^23^, where combined loss of these P4-ATPases abolished plasma membrane PS flippase activity but without constitutive PS exposure.

**Figure 4:**
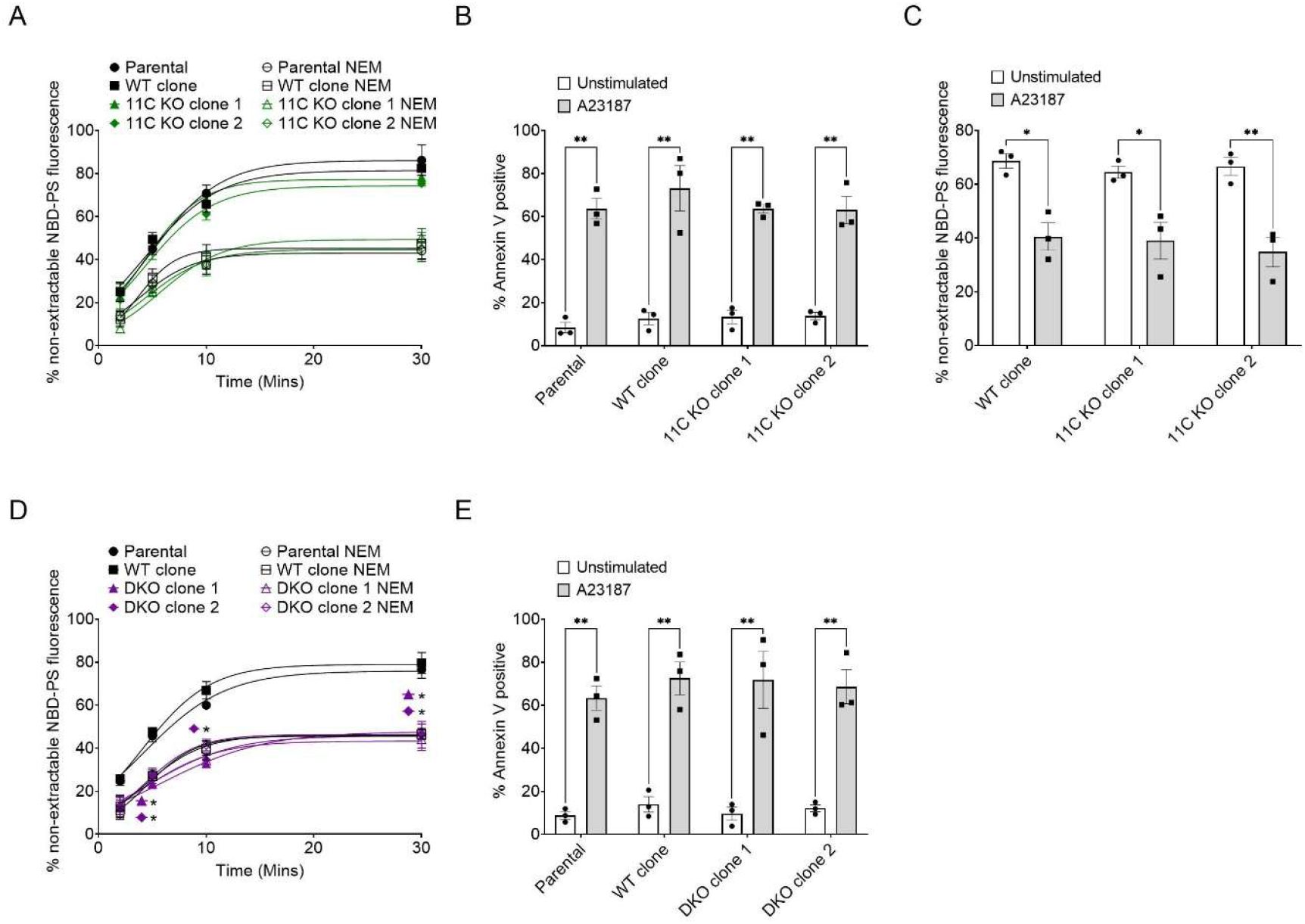
There is full loss of NEM-sensitive flippase activity in DKO megakaryocytes. **A&D**. iQOLG1.1A FoP megakaryocytes were incubated with NEM (5mM: 15 minutes) or vehicle control and incubated with NBD-PS (5μM: 37°C) before sampling into HBS± 2% BSA. **B&E**. FoP megakaryocytes were stimulated with A23187 (10μM: 10 minutes: with 2mM Ca^2+^) and stained with Annexin V to detect PS exposure. **C**. FoP megakaryocytes were stimulated with A23187 (10μM: 10 minutes: with 2mM Ca^2+^) and incubated with NBD-PS (5μM: 37°C: 30 minutes) before sampling into HBS± 2% BSA. Data analysed by two-way ANOVA followed by Dunnett’s multiple comparisons compared with untreated parental FoP MKs (A&D) or unstimulated FoP MKs (B,C,E) (*p<0.05, **p<0.01, ***p<0.001). Mean ± SEM (N=3).

With ATP11A and ATP11C implicated in human megakaryocyte and therefore platelet PS flippase activity, they should now be considered as possible targets for antithrombotic therapies. Maintenance of flippase activity in procoagulant platelets is sufficient to reduce PS exposure and thrombin generation *in vitro*^*15*^. Identification of compounds capable of maintaining the activity of either ATP11A or ATP11C in the presence of a high cytosolic Ca^2+^ signal could therefore have important clinical applications.

In this study, we identify ATP11A and ATP11C in FoP megakaryocytes as acting redundantly to internalise plasma membrane PS. The use of 2 clones each across two independent human iPSC lines shows that these phenotypes are independent of clonal and genetic variability. This study implicates ATP11A and ATP11C as the most likely candidates for the human platelet PS flippase.

## References

1. World Health Organisation. The top 10 causes of death. https://www.who.int/news-room/fact-sheets/detail/the-top-10-causes-of-death (2024).

2. Shah, A., Kerner, V., Stanworth, S. J. & Agarwal, S. Major haemorrhage: past, present and future. Anaesthesia 78, 93–104 (2023).

3. Chauke, L., Bhoora, S. & Ngene, N. C. Postpartum haemorrhage - an insurmountable problem? Case Rep Womens Health 37, e00482 (2023).

4. Munnix, I. C. A. et al. Segregation of Platelet Aggregatory and Procoagulant Microdomains in Thrombus Formation. Arterioscler Thromb Vasc Biol 27, 2484–2490 (2007).

5. Wolfs, J. L. N. et al. Activated scramblase and inhibited aminophospholipid translocase cause phosphatidylserine exposure in a distinct platelet fraction. Cellular and Molecular Life Sciences 62, 1514–1525 (2005).

6. Suzuki, J., Umeda, M., Sims, P. J. & Nagata, S. Calcium-dependent phospholipid scrambling by TMEM16F. Nature 468, 834–838 (2010).

7. Abbasian, N., Millington-Burgess, S. L., Chabra, S.Malcor, J.-D. & Harper, M. T. Supramaximal calcium signaling triggers procoagulant platelet formation. Blood Adv 4, 154–164 (2020).

8. Jobe, S. M. et al. Critical role for the mitochondrial permeability transition pore and cyclophilin D in platelet activation and thrombosis. Blood 111, 1257–1265 (2008).

9. Millington-Burgess, S. L. & Harper, M. T. Gene of the issue: ANO6 and Scott Syndrome. Platelets 31, 964–967 (2020).

10. Yang, H. et al. TMEM16F Forms a Ca2+-Activated Cation Channel Required for Lipid Scrambling in Platelets during Blood Coagulation. Cell 151, 111–122 (2012).

11. Fujii, T., Sakata, A., Nishimura, S., Eto, K. & Nagata, S. TMEM16F is required for phosphatidylserine exposure and microparticle release in activated mouse platelets. Proceedings of the National Academy of Sciences 112, 12800–12805 (2015).

12. Baig, A. A. et al. TMEM16F-Mediated Platelet Membrane Phospholipid Scrambling Is Critical for Hemostasis and Thrombosis but not Thromboinflammation in Mice—Brief Report. Arterioscler Thromb Vasc Biol 36, 2152–2157 (2016).

13. Tilly, R. H. J., Senden, J. M. G., Comfurius, P., Bevers, E. M. & Zwaal, R. F. A. Increased aminophospholipid translocase activity in human platelets during secretion. Biochimica et Biophysica Acta (BBA) - Biomembranes 1029, 188–190 (1990).

14. Basse, F., Gaffet, P., Rendu, F. & Bienvenue, A. Translocation of spin-labeled phospholipids through plasma membrane during thrombin- and ionophore A23187-induced platelet activation. Biochemistry 32, 2337–2344 (1993).

15. Millington-Burgess, S. L. & Harper, M. T. Maintaining flippase activity in procoagulant platelets is a novel approach to reducing thrombin generation. Journal of Thrombosis and Haemostasis https://doi.org/10.1111/jth.15641 (2022) xdoi:10.1111/jth.15641.

16. Sune, A., Bette-Bobillo, P., Bienvenue, A., Fellmann, P. & Devaux, P. F. Selective outside-inside translocation of aminophospholipids in human platelets. Biochemistry 26, 2972– 2978 (1987).

17. Moriyama, Y. & Nelson, N. Purification and properties of a vanadate- and N-ethylmaleimide-sensitive ATPase from chromaffin granule membranes. Journal of Biological Chemistry 263, 8521–8527 (1988).

18. Coleman, J. A., Quazi, F. & Molday, R. S. Mammalian P4-ATPases and ABC transporters and their role in phospholipid transport. Biochimica et Biophysica Acta (BBA) - Molecular and Cell Biology of Lipids 1831, 555–574 (2013).

19. van der Velden, L. M. et al. Heteromeric Interactions Required for Abundance and Subcellular Localization of Human CDC50 Proteins and Class 1 P4-ATPases. Journal of Biological Chemistry 285, 40088–40096 (2010).

20. Segawa, K., Kurata, S. & Nagata, S. The CDC50A extracellular domain is required for forming a functional complex with and chaperoning phospholipid flippases to the plasma membrane. Journal of Biological Chemistry 293, 2172–2182 (2018).

21. Segawa, K. et al. Caspase-mediated cleavage of phospholipid flippase for apoptotic phosphatidylserine exposure. Science (1979) 344, 1164–1168 (2014).

22. Wang, W. et al. Mobilizing phospholipids on tumor plasma membrane implicates phosphatidylserine externalization blockade for cancer immunotherapy. Cell Rep 41, 111582 (2022).

23. Miyata, Y., Yamada, K., Nagata, S. & Segawa, K. Two types of type IV P-type ATPases independently re-establish the asymmetrical distribution of phosphatidylserine in plasma membranes. Journal of Biological Chemistry 102527 (2022) doi:10.1016/j.jbc.2022.102527.

24. Takatsu, H. et al. ATP9B, a P4-ATPase (a Putative Aminophospholipid Translocase), Localizes to the trans-Golgi Network in a CDC50 Protein-independent Manner. Journal of Biological Chemistry 286, 38159–38167 (2011).

25. Takatsu, H. et al. Phospholipid Flippase Activities and Substrate Specificities of Human Type IV P-type ATPases Localized to the Plasma Membrane. Journal of Biological Chemistry 289, 33543–33556 (2014).

26. Segawa, K., Kurata, S. & Nagata, S. Human Type IV P-type ATPases That Work as Plasma Membrane Phospholipid Flippases and Their Regulation by Caspase and Calcium. Journal of Biological Chemistry 291, 762–772 (2016).

27. Gyobu, S., Ishihara, K., Suzuki, J., Segawa, K. & Nagata, S. Characterization of the scrambling domain of the TMEM16 family. Proceedings of the National Academy of Sciences 114, 6274–6279 (2017).

28. Segawa, K. et al. Phospholipid flippases enable precursor B cells to flee engulfment by macrophages. Proceedings of the National Academy of Sciences 115, 12212–12217 (2018).

29. Ochiai, Y., Suzuki, C., Segawa, K., Uchiyama, Y. & Nagata, S. Inefficient development of syncytiotrophoblasts in the Atp11a -deficient mouse placenta. Proceedings of the National Academy of Sciences 119, (2022).

30. Arashiki, N. et al. ATP11C Encodes a Major Flippase in Human Erythrocyte and Its Genetic Defect Causes Congenital Non-Spherocytic Hemolytic Anemia. Blood 126, 2131–2131 (2015).

31. Arashiki, N. et al. ATP11C is a major flippase in human erythrocytes and its defect causes congenital hemolytic anemia. Haematologica 101, 559–565 (2016).

32. Mansour-Hendili, L. et al. Exome sequencing for diagnosis of congenital hemolytic anemia. Orphanet J Rare Dis 15, 180 (2020).

33. van Dijk, M. J. et al. A novel missense variant in ATP11C is associated with reduced red blood cell phosphatidylserine flippase activity and mild hereditary hemolytic anemia. Am J Hematol 98, 1877–1887 (2023).

34. Fermo, E. et al. Not so Rare, Not so Mild Disease: Defects of Flippase Activity Due to ATP11C Mutations. Description of Three New Cases. Blood 144, 2465–2465 (2024).

35. Moreau, T. et al. Large-scale production of megakaryocytes from human pluripotent stem cells by chemically defined forward programming. Nat Commun 7, 11208 (2016).

36. Evans, A. L. et al. Transfer to the clinic: refining forward programming of hPSCs to megakaryocytes for platelet production in bioreactors. Blood Adv 5, 1977–1990 (2021).

37. Putz, M. V., Lacrama, A. M. & Ostafe, V. Introducing logistic enzyme kinetics. Journal of Optoelectronics and Advanced Materials 9, 2910–2916 (2007).

38. Daleke, D. L., Cornely-Moss, K., Lyles, J., Smith, C. M. & Zimmerman, M. Identification and Characterization of a Candidate Phosphatidylserine-Transporting ATPase. Ann N Y Acad Sci 671, 468–470 (1992).

