## Supplementary figures for "ATP11A and ATP11C are plasma membrane phosphatidylserine flippases in *in vitro* human megakaryocytes"

**Supplementary material**

**Supplementary Table 1 – Components of megakaryocyte media (AMK).**

| Cell culture component | Volume/Concentration | Supplier | Catalogue no. |
|---|---|---|---|
| Basal media IMDM without phenol red | 500ml | Thermofisher | 21056023 |
| Chemically defined lipid concentrate | 5ml | Thermofisher | 11905031 |
| 100x Insulin-Transferrin-Selenium | 5ml | Thermofisher | 41400045 |
| 2-Mercaptoethanol | 0.5ml | Thermofisher | 10306050 |
| Bovine Albumin Serum (30%) | 8.4ml | Lab-Tech | SA-296 |

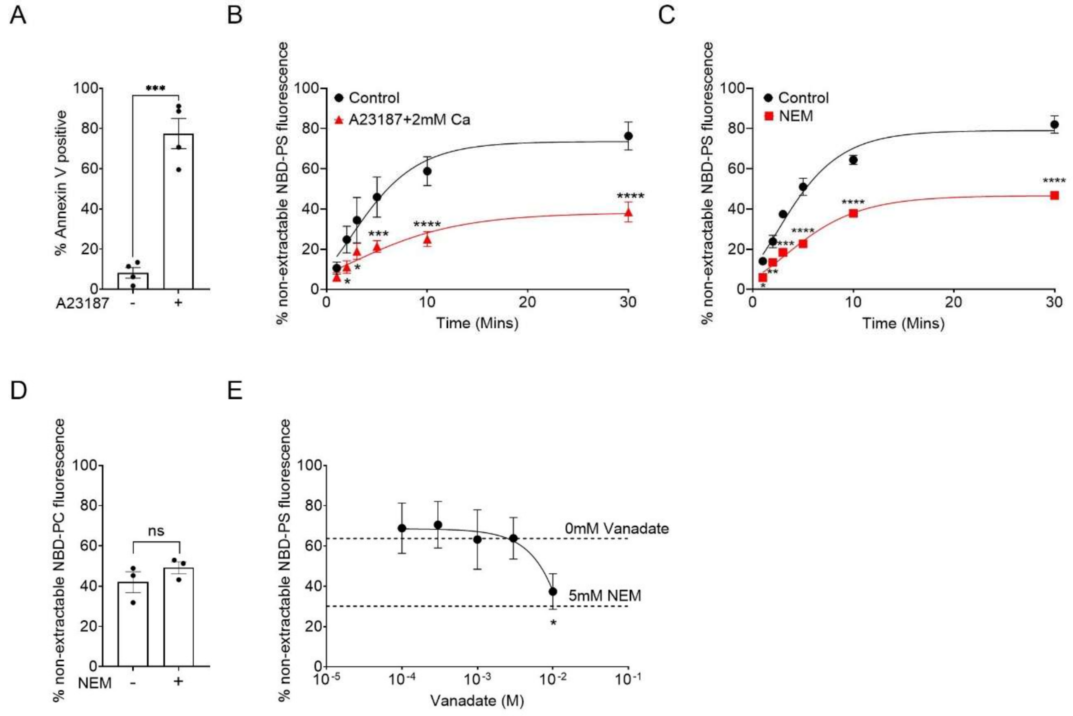


**Supplementary Figure 1: Flippase activity in FoP megakaryocytes from hiPSC line 2. A.** FoP megakaryocytes (5x$10^5$/ml) from iLIPSC-GR1.1.11A were stimulated with A23187 (10μM: 10 minutes; with 2mM $CaCl_2$). Cells were then stained with Annexin V to detect PS exposure. **B-E.** FoP megakaryocytes were treated as indicated with A23187, NEM (5μM: 15 minutes), vanadate (various concentrations: 15 minutes), or vehicle controls, and incubated with NBD-PS (**B,C,E** 5μM: 37°C) or NBD-PC (**D** 5μM: 37°C: 30 minutes) for the indicated times before sampling into HBS± 2% BSA. Data analysed by paired Students T test **(A&D)**, one-way ANOVA **(E)** or two-way ANOVA **(B&C)** followed by Dunnett's multiple comparisons (*p<0.05, **p<0.01, ***p<0.001, ****p<0.0001). Mean ± SEM (N=3-4)

**Supplementary Table 2: Summary of mutations introduced into CDC50A and the size of the truncated product.**

| Cell line | Clone | Mutation CDC50A |
|---|---|---|
| iQOLG1.1A | CDC50A-KO Clone 1 | -7 (180 AA) |
| | CDC50A-KO Clone 2 | +1 (143AA) |
| iLIPSC-GR1.1.11A | CDC50A-KO Clone 1 | +1 (143 AA) |
| | CDC50A-KO Clone 2 | -7 (180AA) |

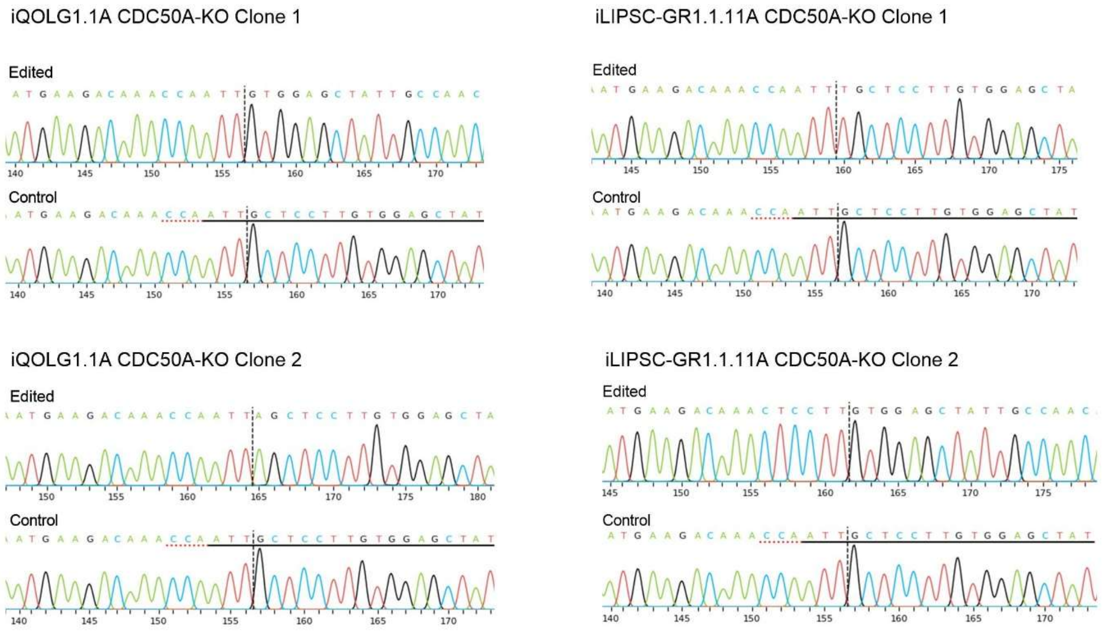


**Supplementary Figure 2: Sequencing data for iQOLG1.1A and iLIPSC-GR1.1.11A CDC50A-KO clones.** Edited and control Sanger sequencing chromatograms from the Inference of CRISPR Edits (ICE) online tool for iQOLG1.1A and iLIPSC-GR1.1.11A CDC50A-KO clones.

**Supplementary Table 3: Summary of mutations introduced into ATP11A and ATP11C and the size of the truncated product.**

| Cell line | Clone | Mutation ATP11A | Mutation ATP11C |
|---|---|---|---|
| iQOLG1.1A | ATP11A-KO Clone 1 | +1 (66 AA) | WT (1132 AA) |
| | ATP11A-KO Clone 2 | +1 (66 AA), -11 (62 AA) | WT (1132 AA) |
| | ATP11C-KO Clone 1 | WT (1134 AA) | -4 (247 AA) |
| | ATP11C-KO Clone 2 | WT (1134 AA) | +1 (181 AA) |
| | DKO Clone 1 | +1 (66 AA) | +1 (181AA) |
| | DKO Clone 2 | +1 (66 AA) | -7 (246 AA) |
| iLIPSC-GR1.1.11A | ATP11A-KO Clone 1 | +1 (66 AA), -16 (66 AA) | WT (1132 AA) |
| | ATP11A-KO Clone 2 | +1 (66 AA) | WT (1132 AA) |
| | ATP11C-KO Clone 1 | WT (1134 AA) | -1 (248 AA) |
| | ATP11C-KO Clone 2 | WT (1134 AA) | +1 (181 AA) |
| | DKO Clone 1 | +1 (66 AA), -2 (65AA) | +1 (181 AA) |
| | DKO Clone 2 | +1 (66 AA) | +1 (181 AA) |

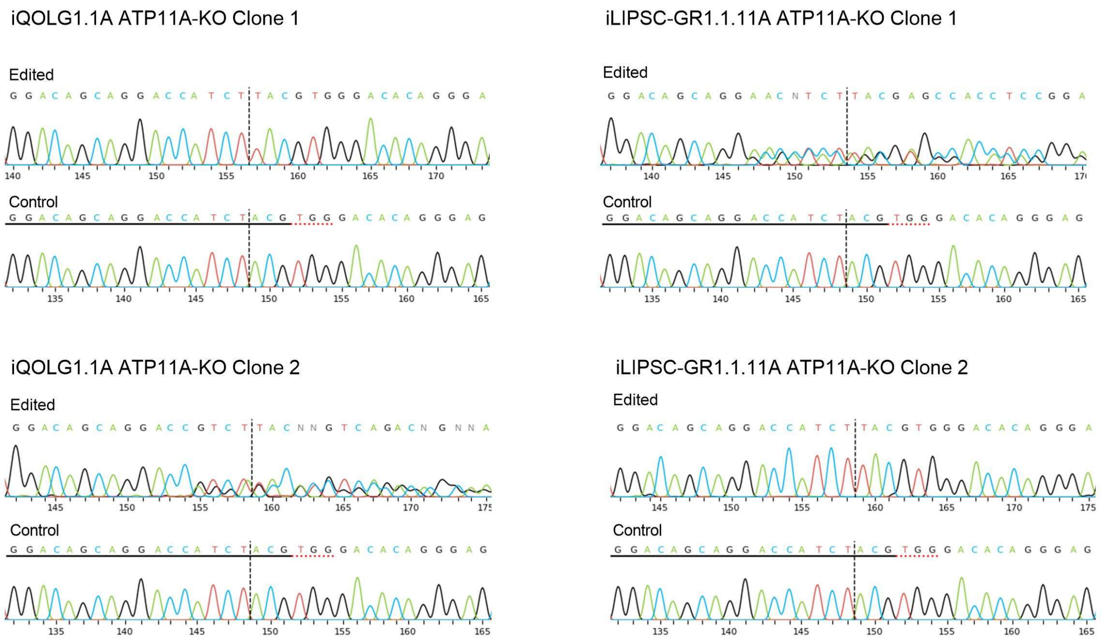


**Supplementary Figure 3: Sequencing data for iQOLG1.1A and iLIPSC-GR1.1.11A ATP11A-KO clones.** Edited and control Sanger sequencing chromatograms from the ICE online tool for iQOLG1.1A and iLIPSC-GR1.1.11A ATP11A-KO clones.

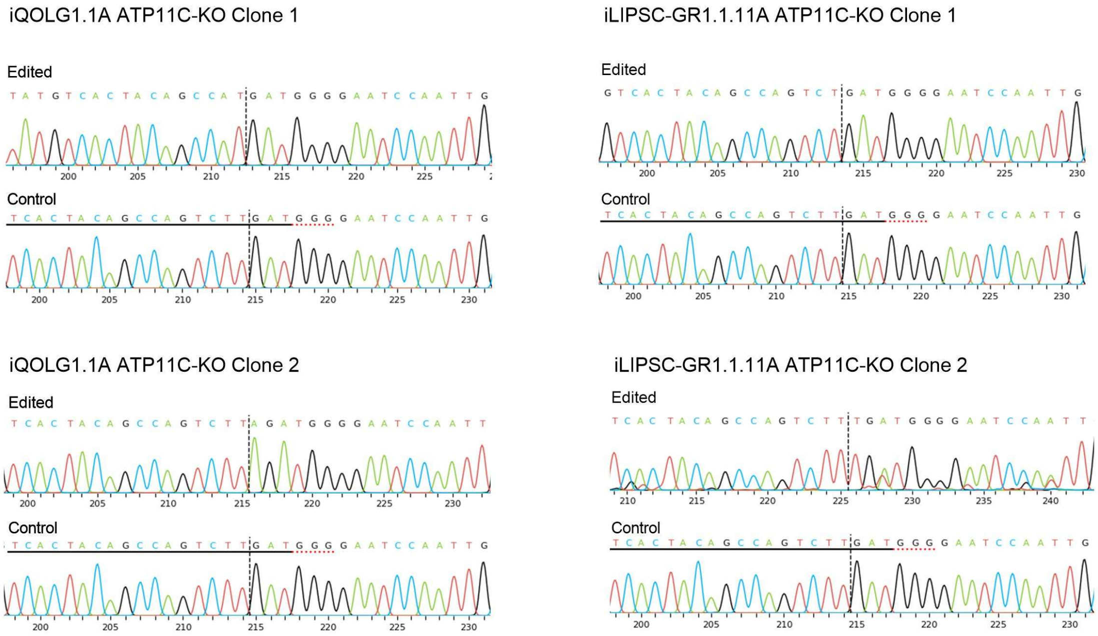


**Supplementary Figure 4: Sequencing data for iQOLG1.1A and iLIPSC-GR1.1.11A ATP11C-KO clones.** Edited and control Sanger sequencing chromatograms from the ICE online tool for iQOLG1.1A and iLIPSC-GR1.1.11A ATP11C-KO clones.

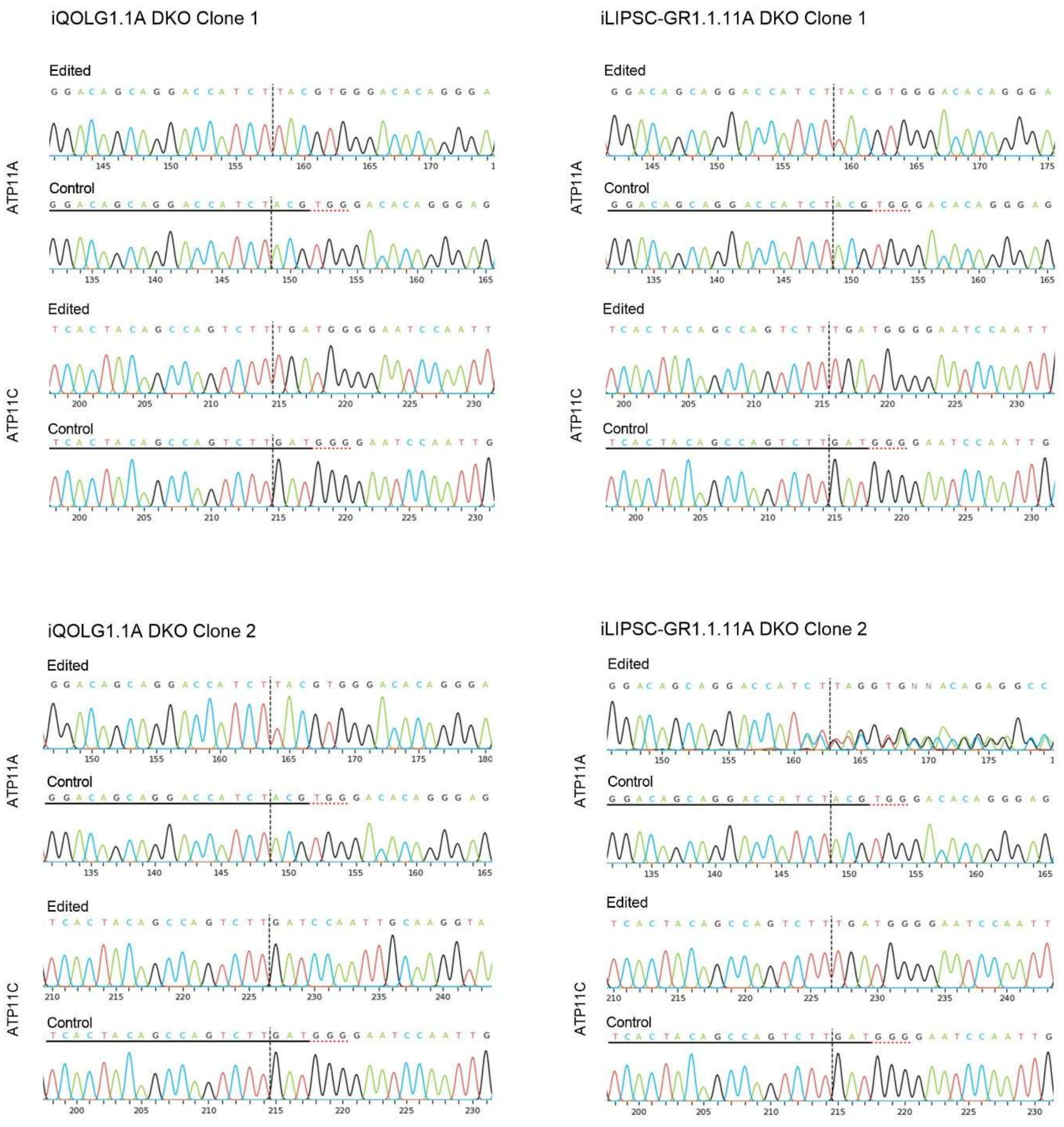


**Supplementary Figure 5: Sequencing data for iQOLG1.1A and iLIPSC-GR1.1.11A DKO clones.** Edited and control Sanger sequencing chromatograms from the ICE online tool for iQOLG1.1A and iLIPSC-GR1.1.11A DKO clones.

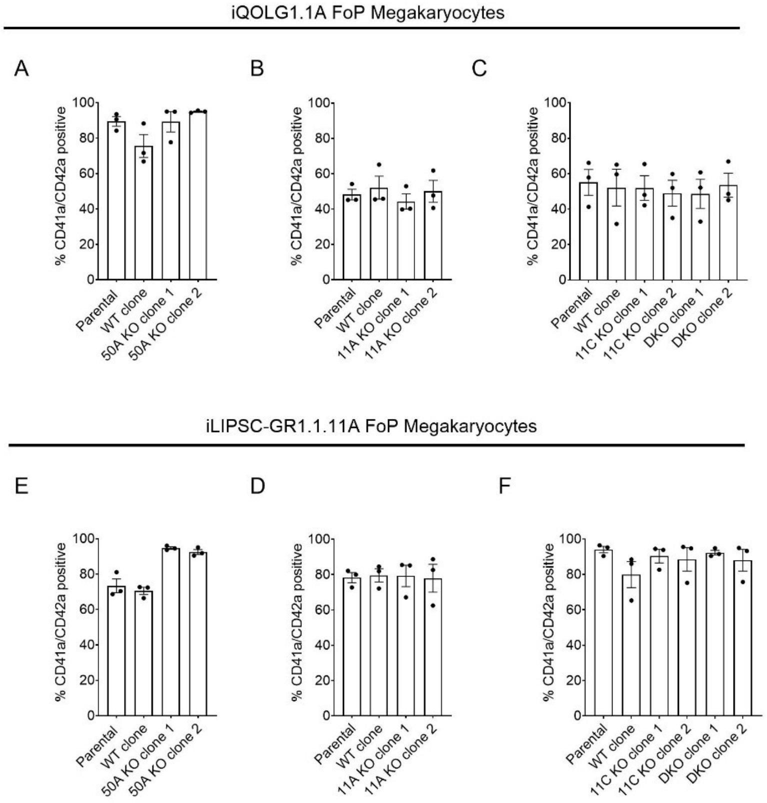


**Supplementary Figure 6: The percentage of viable Day 20 cells that are positive for CD41a and CD42a was not significantly different when compared with the parental population.** **A-D** Day 20 cells from iQOLG1.1A (**A-C**) and iLIPSC-GR1.1.11A (**D-F**) were stained for CD41a (1 in 100) and CD42a (1 in 200) using DAPI as a marker of viability (1 in 200). Mean ± SEM (N=3).

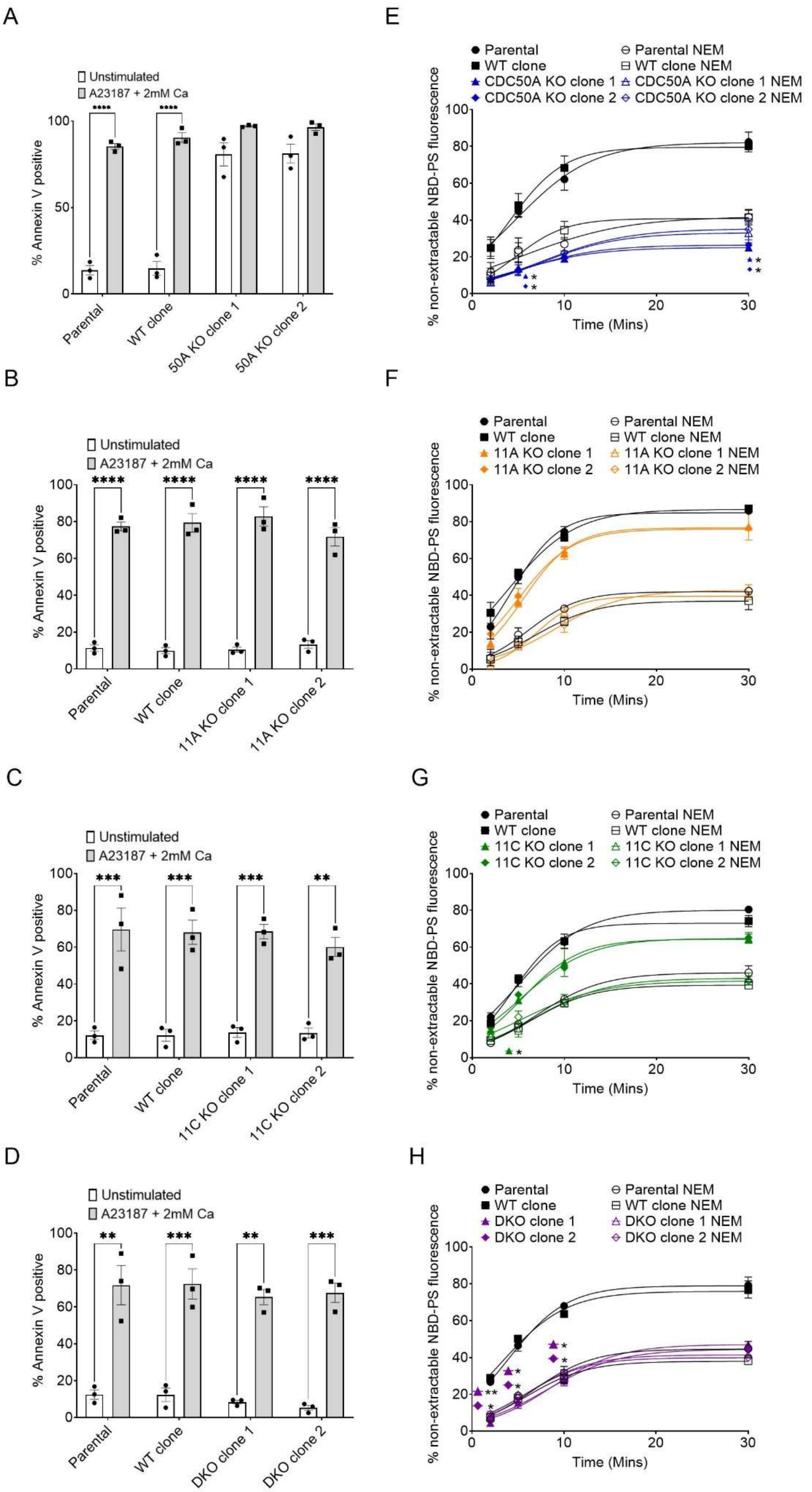

A
Unstimulated
A23187 + 2mM Ca
% Annexin V positive
Parental
WT clone
50A KO clone 1
50A KO clone 2
B
Unstimulated
A23187 + 2mM Ca
% Annexin V positive
Parental
WT clone
11A KO clone 1
11A KO clone 2
C
Unstimulated
A23187 + 2mM Ca
% Annexin V positive
Parental
WT clone
11C KO clone 1
11C KO clone 2
D
Unstimulated
A23187 + 2mM Ca
% Annexin V positive
Parental
WT clone
DKO clone 1
DKO clone 2
E
Parental
WT clone
CDC50A KO clone 1
CDC50A KO clone 2
Parental NEM
WT clone NEM
CDC50A KO clone 1 NEM
CDC50A KO clone 2 NEM
% non-extractable NBD-PS fluorescence
Time (Mins)
F
Parental
WT clone
11A KO clone 1
11A KO clone 2
Parental NEM
WT clone NEM
11A KO clone 1 NEM
11A KO clone 2 NEM
% non-extractable NBD-PS fluorescence
Time (Mins)
G
Parental
WT clone
11C KO clone 1
11C KO clone 2
Parental NEM
WT clone NEM
11C KO clone 1 NEM
11C KO clone 2 NEM
% non-extractable NBD-PS fluorescence
Time (Mins)
H
Parental
WT clone
DKO clone 1
DKO clone 2
Parental NEM
WT clone NEM
DKO clone 1 NEM
DKO clone 2 NEM
% non-extractable NBD-PS fluorescence
Time (Mins)

**Supplementary Figure 7: Phenotypic analysis of iLIPSC-GR1.1.11A ATP11A-KO, ATP11C-KO and DKO FoP megakaryocytes. A-D**. iLIPSC-GR1.1.11A FoP megakaryocytes were stimulated with A23187 (10µM: 10 minutes: with 2mM $Ca^{2+}$) and stained with Annexin V to detect PS exposure. **E-H.** FoP megakaryocytes were incubated with NEM (5mM: 15 minutes) or vehicle control and incubated with NBD-PS (5µM: 37°C) before sampling into HBS± 2% BSA. Data analysed by two-way ANOVA followed by Dunnett's multiple comparisons compared with unstimulated FoP MKs (A-D) or untreated parental FoP MKs (E-H) (*p<0.05, **p<0.01, ***p<0.001, ****p<0.0001). Mean ± SEM (N=3).
